# Species delimitation and phylogeny estimation under the multispecies coalescent

**DOI:** 10.1101/010199

**Authors:** Graham Jones

## Abstract

This article describes a Bayesian method for inferring both species delimitations and species trees under the multispecies coalescent model using DNA sequences from multiple loci. The focus here is on species delimitation with no a priori assignment of individuals to species, and no guide tree. The method uses a new model for the population sizes along the branches of the species tree, and three new operators for sampling from the posterior using the Markov chain Monte Carlo (MCMC) algorithm. The correctness of the moves is demonstrated both by proofs and by tests of the implementation. Current practice, using a pipeline approach to species delimitation under the multispecies coalescent, has been shown to have major problems on simulated data [10]. The same simulated data set is used to demonstrate the accuracy and efficiency of the present method. The method is implemented in a package called STACEY for BEAST2.

## 1 Introduction

Species delimitation is the problem of assigning a number of individual organisms to one or more species. The word ‘delimitation’ is also used to refer to a particular assignment or clustering of the individuals into groups or clusters. There are many approaches to this important problem and this article concentrates on the use of genetic data to infer such a clustering. The basic idea can be understood by considering two gene copies sampled at present time. Tracing their history back in time, they can coalesce within a group of interbreeding organisms according to a coalescent model, whereas between different groups, they cannot coalesce until the groups have merged. This idea is made precise by the multispecies coalescent model [15, 13, 2, 3].

There are two main types of ‘noise’ which interfere with inference of delimitation and phylogeny: mutational variance and incomplete lineage sorting. Mutational variance is a problem because the organisms are often closely related with few mutations separating them. This means there is a large amount of uncertainty about coalescence times. Incomplete lineage sorting refers to gene copies which fail to coalesce until after the (single) species to which they both belong has merged with another species.The degree of incomplete lineage sorting depends the effective population size along a branch, divided by the branch length measured in generations.

### 1.1 Previous work

Over the past ten years, the multispecies coalescent model has become the standard approach for species tree estimation using sequences from multiple loci. It accounts for incomplete lineage sorting, a ubiquitous source of discord between gene trees. More recently, it has been used for species delimitation, in BPP [16, 14, 17] and DISSECT [9]. More details and discussion of alternative approaches can be found in [9], [10], and [18].

DISSECT uses a prior for the species tree in which the usual birth-death model is replaced by one which incorporates a spike near zero in the density for node heights, a ‘birth-death-collapse’ model. This is a computational approximation to a model in which the dimensionality of the parameter space changes as the number of species changes.

In [10] sequences were simulated under the multispecies coalescent model, and then analyzed the data using a standard ‘pipeline’ method using Structurama [12], [8], *BEAST [5], and BPP [14]. The analysis showed there were major problems with the method.

### 1.2 Overview of the present method

The method presented here replaces this pipeline with a single analysis. It is implemented as a package called STACEY (Species Tree And Classification Estimation, Yarely) for BEAST2 [1]. STACEY is aimed mainly at species delimitation, but can also be used as an alternative to *BEAST [5]. It incorporates a new model for the populations along the branches of the SMC-tree, and three new MCMC moves for exploring the posterior when the multispecies coalescent model is assumed. It uses the same birth-death-collapse model as DISSECT.

The multispecies coalescent model requires a model for the populations along the branches of the SMC-tree. The simplest option is to assume that the population in all branches is identical and constant along each branch. Another option, as used in *BEAST, is to introduce one or more population parameters for each branch and sample these using the Markov chain Monte Carlo (MCMC) algorithm. Here is assumed that each branch in has a population parameter which is constant along the branch, and that these parameters are independent and identically distributed. Instead of sampling these parameters, they are integrated out. The method caters for variation among branches, and is similar to the ‘piecewise constant’ option in *BEAST but does not allow individual populations to be estimated. The hope is that this simplification makes the posterior easier to sample from.

To achieve this sampling, operators with the right statistical properties (MCMC moves) are needed. Their design is important for the efficiency of the method. The moves described here were designed with species delimitation in mind, although all three moves are also applicable to species tree estimation with a fixed species delimitation. Species delimitation presents a difficult challenge for the MCMC algorithm, since the MCMC moves must be capable of efficiently exploring all possible delimitations, and for each delimitation, all the usual parameters. In the multi-species coalescent model, there is one species tree and one or more gene trees. Each gene tree must ‘fit inside’ the species tree. A change to the species tree or to a gene tree may result in an incompatibility between the species tree and one or more gene trees. If an MCMC move makes such a change it must be rejected, and if such rejections are common the move will be inefficient. The three moves described here preserve compatibility between the species tree and the gene trees.

The first MCMC move, called NodesNudge, changes the height of a node in a SMC-tree, and changes the height of certain ‘nearby’ nodes in the gene trees. It does this in a way that leaves the all tree topologies unchanged, and preserves the compatibility of the gene trees with the SMC-tree. It is a subtle move, in that it typically changes the node heights by a small amount, but it appears to have a large beneficial effect on the convergence, at least on some data sets.

The second move, called CoordinatedPruneRegraft, is a subtree-prune-and-regraft move which makes coordinated topological changes to the species tree and gene trees. The CoordinatedPruneRegraft move can be seen as an extension of the nearest neighbor interchange (NNI) move described in [17], which makes a coordinated set of fixed node height NNI moves to the species tree and to the gene trees. When viewed this way, the CoordinatedPruneRegraft extends the NNI move to the more general subtree-prune-and-regraft move. It can also be seen as an extension to the ‘Fixed Nodeheight Prune and Regraft’ (FNPR) as described in [6]. The FNPR move changes the topology of a single tree, whereas the move described here makes a coordinated set of FNPR moves to the species tree and to the gene trees in order to maintain compatibility between the trees.

The third move, called FocusedScaler, scales node heights whilst preserving topologies. The scaling is ‘focused’ on a node in the species tree. This node is scaled by the largest amount. The further away a node is from the focus (in a sense to be made precise later), the less it is affected by the move. Once the relative amount by which each node should be scaled by has been chosen, the maximum range of scaling consistent with compatibility is found. The actual scaling is then chosen from this range.

The population model is described first, followed by the MCMC moves. Two sets of tests on simulated data are then described. Firstly, the method is tested for correctness by sampling from prior distributions in cases where some analytic results are available. Finally, the simulated data set of [10] is re-analyzed.

## 2 Conventions and notation

All trees are rooted and binary. Time is measured backwards from zero at present, and all tree nodes have a time, referred to as a node height. A tree topology should be understood as a labeled topology, that is, it includes the assignment of labels to tips. In the context of species delimitation using STACEY, the species tree has tips which represent **minimal clusters** of individuals [9]. These minimal clusters may be merged but not split to form potential species. At its most flexible, there is just one individual in each minimal cluster, so the possible number of species ranges from one to the number of individuals. Thus ‘species tree’ is not a good name for this tree, and instead I will refer to it a the **SMC-tree**, as a shorthand for ‘species or minimal clusters tree’.

Lower case letters are used for gene tree nodes, and upper case for SMC-tree nodes and in situations where the type of tree does not matter. For either type of node *X*, its parent is denoted by anc(*X*) and its node height by *t*(*X*). The branch that leads from anc(*X*) to *X* is referred to as ‘the branch *X*’. The ‘subtree of *X*’ contains *X*, all its descendants, and the branch *X*, but not the node anc(*X*) which is the origin of the subtree.

For a node *X* in the SMC-tree, let *I*(*X*) denote the set of minimal clusters belonging to *X* (that is, assigned to a tip node which is a descendant of *X*). For a node *x* in a gene tree, let *I*(*x*) denote the set of the minimal clusters which yielded a sequence belonging to *x*. Furthermore, if *X* is not a tip, let *R*(*X*) and *L*(*X*) denote the set of minimal clusters belonging to the two children of *X*. Note that *I*(*X*) = *L*(*X*) ∪ *R*(*X*) for both SMC-tree nodes and gene tree nodes, so they can be calculated recursively from the tips, and all these sets are unions of minimal clusters. In the SMC-tree the unions are disjoint, and a node is uniquely identified by its set of minimal clusters. In the gene tree case, neither of these is true in general. However, the set *I*(*x*) and height *t*(*x*) for a gene tree node *x* are enough to assign *x* to a unique branch in the SMC-tree, as follows. If the SMC-tree is cut across at height *t*(*x*), this will intersect some branches *X*_1_*, X*_2_*, …, X_n_* say. All the *I*(*X*_*i*_) are pairwise disjoint, and *I*(*x*) cannot intersect more than one of them non-trivially or the gene tree would be incompatible with the SMC-tree. Thus *I*(*x*) ⊂ *I*(*X*_*i*_) for some *i* thus identifying the branch *X*_*i*_ as the one which contains *x*.

The notion that part of a gene tree is ‘inside’ a branch of a SMC-tree is intuitively obvious from diagrams, but a formal definition is required for algorithms and proofs. Suppose *X* is a node in the SMC-tree and *x* is a gene tree node and *t* ∈ [*t*(*x*)*, t*(anc(*x*))]. Then the point (*x, t*) is **inside** the branch *X* if *I*(*x*) ⊂ *I*(*X*) and *t* ∈ [*t*(*x*)*, t*(anc(*X*))].

We also formally define the notion of compatibility. Firstly, if *A* is a set of minimal clusters, and *X* is a node in the SMC-tree, we say that *A* **straddles** *X* if *X* is not a tip, *A ∩ L*(*X*) ≠ ø, and *A ∩ R*(*X*) ≠ ø. If *X* is a node in the SMC-tree and *x* is a gene tree node, then the pair (*X, x*) is **compatible** if *t*(*x*) ≥ *t*(*X*) or *I*(*x*) does not straddle *X*. A gene tree is compatible with the SMC-tree if every pair of nodes (*X, x*) is compatible.

We define a pair of nodes (*X, x*) with *X* in the SMC-tree and *x* in a gene tree to be **hitched** if *I*(*x*) straddles *X*, but neither *L*(*x*) nor *R*(*x*) straddle *X*. See Figure 1 for an example. The hitched nodes are the minimal set of nodes that need to be checked for compatibility to ensure the SMC-tree and the gene tree are compatible. See Figure 1.

**Figure 1:**
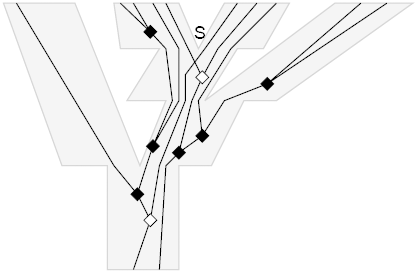
Example of hitched nodes. The SMC-tree is pale gray. A gene tree is shown inside it. Gene tree nodes which are hitched to the SMC-tree node *S* are shown as white diamonds, and other nodes as black diamonds.

## 3 The population model

Consider a single gene and a single branch. The coalescent model of Kingman (see Chapters 26-28 of [4]) is assumed. The probability density for the coalescent times takes the following form (simplified from equation (3), p572, of [5]):

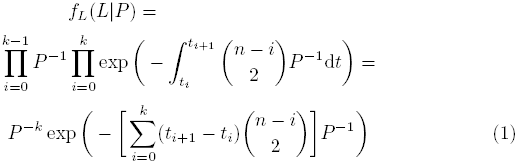

where *L* is the lineage history of a gene tree within a single branch, and *P* is the effective number of gene copies in the population for this branch, which is assumed constant along the branch in this paper. Thus *P* is the expected number of generations for a pair of gene copies to coalesce. The lineage history *L* consists of the number *n* of lineages at the tipward end of the branch, the number *k* of coalescences within the branch, plus the times (*t*_0_ *< t*_1_*, …t_k_ < t*_*k*+1_) where *t*_0_ is the node height at the tipward end, *t*_*k*+1_ is the node height at the rootward end, and (*t*_1_*, …t_k_*) are the coalescence times within the branch. Between *t*_*i*_ and *t*_*i*+1_ there are *n − i* lineages. The complete multispecies coalescent probability density is the double product, over genes and over branches, of terms like this.

As usual, we convert *P* into substitution units by multiplying by the mutation rate measured in substitutions per site per generation. Denote the effective population in branch *b* by *N*_*b*_ and the mutation rate by *μ_b_*. The effective number of gene copies is obtained from *N*_*b*_ by multiplying by a factor *p*_*j*_ (sometimes called the ‘ploidy’) for gene *j*. This *p*_*j*_ depends on the type of gene involved, and is 2 for the common case of autosomal nuclear genes in diploid species. Exceptions include genes from sex chromosomes and organelles. For gene *j* in branch *b*, we thus need to replace *P* by *p*_*j*_*N*_*b*_*μ*_*b*_ in equation (1).

To write down the full expression, some more notation is needed. The branches in the SMC-tree are indexed by *b*. A sum or product over *b* should be understood as being over all branches. Note that this includes the root, so that all gene lineages eventually coalesce. The number of branches is *B*. Set *θ_b_* = *N*_*b*_*μ*_*b*_. The vector (*θ*_1_*, θ*_2_*, …, θ_B_*) is denoted by Θ. The genes are indexed by *j*. A sum or product over *j* should be understood as being over all genes. The number of coalescences of gene *j* within branch *b* is denoted by *k*_*jb*_. The number of lineages in gene tree *j* at the tipward end of branch *b* is denoted by *n*_*jb*_. The number of lineages in gene tree *j* at the rootward end of branch *b* is thus *n*_*jb*_ − *k*_*jb*_. The time interval between the tipward and rootward branch *b* is divided into *k*_*jb*_ + 1 intervals by the coalescent times of gene *j*. These *k*_*jb*_ + 1 intervals are denoted by *c*_*jbi*_ (0 *≤ i ≤ k_jb_*). There are *n*_*jb*_ − *i* lineages in gene tree *j*, branch *b* during the time interval *c*_*jbi*_. Let *G* denote all the lineage histories of all the genes in all the branches. The complete multispecies coalescent probability density is

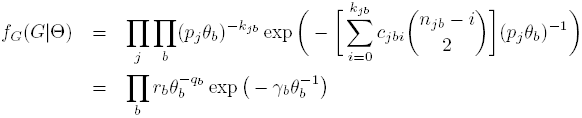

Where

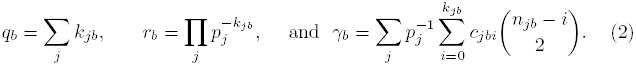

For each *b* this has the form of an unnormalised inverse gamma density for *θ_b_*. The normalised inverse gamma density is

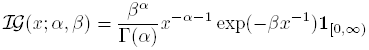

where *α* and *β* are parameters in (0, ∞). If, a priori, the *θ_b_* are assumed independent and are assumed to have an inverse gamma density it is possible to integrate out the *θ_b_* analytically. In fact the prior can be more general than a single inverse gamma density: an overall scaling parameter *σ* can be introduced, together with hyperprior *π_σ_*(*σ*) for it; and a mixture of inverse gamma densities can be used. This mixture takes the form

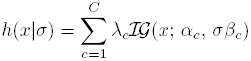

Here *C*, the *λ*_*c*_, the *α*_*c*_, and the *β*_*c*_ (1 ≤ *c* ≤ *C*) are user-chosen values, which are constant for the analysis. The *λ*_*c*_ are positive and sum to one, and the *α*_*c*_ and *β*_*c*_ are arbitrary positive numbers. The density *π*_*σ*_ is also user-chosen and can be any density with support contained in [0, ∞). Each *θ*_*b*_ is then an independent draw from the density *h*. So the joint prior density for Θ and *σ* is

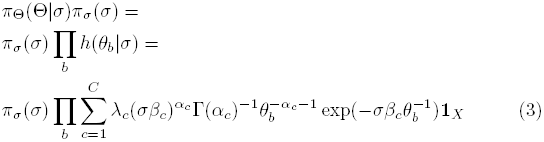

where *X* is the positive orthant in **R**^*B*^.

Then combining (2) and (3), the posterior density for the multispecies coalescent is

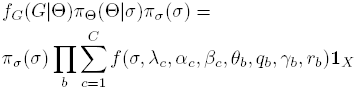

where *f*(*σ*,*λ*_*c*_,*α*_*c*_,*β*_*c*_,*θ*_*b*_,*q*_*b*_,*γ*_*b*_,*r*_*b*_)=

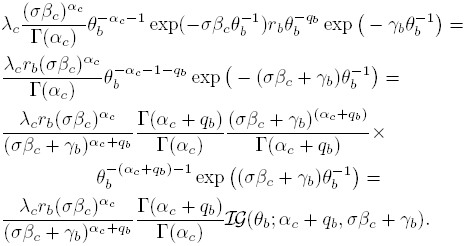

Now Θ can be integrated out from the posterior, using the fact that *𝓘𝓖* integrates to 1 to obtain

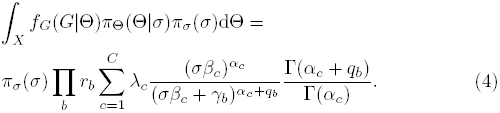

Equations (4) and (2) provide the information needed to implement the method.

## 4 The NodesNudge move

### 4.1 The algorithm

We describe a more general algorithm than the move which is currently implemented, since it may be useful to use variants of the move. It uses the concept of a connected component from graph theory. Given a subset Δ of the nodes in a gene tree, we first remove the nodes not in Δ, then divide what is left into the connected components. Figure 2 illustrates the idea. On the left is a gene tree, in which nodes are shown by solid diamonds if they are in Δ and open diamonds otherwise. On the right, the three connected components in Δ are shown as diamonds and solid lines. For any gene tree node *x* ∈ Δ, let *C*(*x*) denote the connected component in the gene tree to which *x* belongs. Furthermore, define the set *C*^*^(*x*) to be the set of nodes *c* in the gene tree such that anc(*c*) ∈ *C*(*x*) and *c ∉ C*(*x*). Their positions are at the tops of the dotted lines in the right of the figure, and can be thought of as the ‘children’ of *C*(*x*). Finally let *r*(*x*) be the oldest node in *C*(*x*), the root of *C*(*x*). Here is the algorithm:

**Figure 2:**
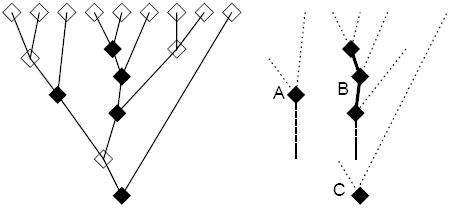
Example of connected components.

1. Choose uniformly and at random any node *S* in the SMC-tree which is not a tip and not the root.
2. Let Δ(*S*) = {*s*_1_*, …, s_n_*} be a set of internal gene tree nodes defined by a criterion which only depends on *S*, and the topologies of the SMC-tree and gene trees.
3. Let *d*_0_ = max_*X*_ *{t*(*X*) : anc(*X*) = *S}*, which is the time of the most ancient of the two child nodes of *S*, and let *u*_0_ = *t*(anc(*S*)).
4. For 1 *≤ i ≤ n*, let

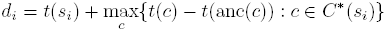

and let *u*_*i*_ = *∞* if *s*_*i*_ is the root, otherwise let *u*_*i*_ = *t*(*s*_*i*_) + *t*(anc(*r*(*s*_*i*_))) *- t*(*r*(*s*_*i*_)).
5. Let

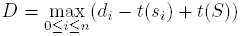

and

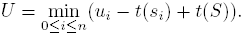
6. Choose a new node height *t′*(*S*) for *S* uniformly in [*D, U*].
7. Change the height of all gene tree nodes in Δ(*S*) by the same amount, that is, by *t′*(*S*) *- t*(*S*).

We formally prove some properties of this move below; here is an informal description of the ideas behind the proofs. Firstly, note that the value *d*_0_ (step 3) can be written as *t*(*S*) + max_*X*_ {*t*(*X*) *- t*(*S*) : anc(*X*) = *S*} which emphasizes the similarity with the other *d*_*i*_ (step 4). Next, note that since *S* is not he root, *u*_0_ is finite, so that [*D, U*] is a finite interval, and step 6 makes sense.

Now we explain the role of the connected components. Returning to Figure 2, note that the minimum length of the dotted edges leaving each connected component determines the maximum amount by which the nodes in the connected component can move forward in time. The oldest node in each connected component usually provides a limit (the length of the dashed line) on how far back in time the connected component can move; the exception is if it is the root node of the gene tree, as in the case of connected component C. The key property of connected components is that the limit of movement back and forwards in time of one connected component is determined by the times of nodes which cannot belong to another connected component. This ensures that the definitions of *D* and *U* are unaffected by the move. Also note that all nodes are moved by the same amount, so the internal structure of *C*(*s*_*i*_) does not change.

### 4.2 Properties

#### Proposition 1

The NodesNudge move preserves all the tree topologies and keeps all branch lengths nonegative.

*Proof.* From step 6, *D ≤ t′*(*S*) ≤ *U*, and from step 5, *d*_0_ ≤ *t′*(*S*) ≤ *u*_0_, and so it follows from step 3 that

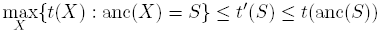

hence the new height for *S* lies between that of *S*’s oldest child and its parent.

Now assume 1 ≤ *i ≤ n*. From step 7, the new height *t′*(*s*_*i*_) is *t*(*s*_*i*_) + *t′*(*S*) *t*(*S*). From steps 5 and 6,

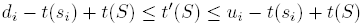

so

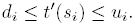

The next step is to show that the definition of *d*_*i*_ and *u*_*i*_ in step 4 preserves the topologies and keeps all branch lengths non-negative in the gene trees. The condition *c* ∈ *C\**(*s*_*i*_) identifies all pairs of nodes (*c,* anc(*c*)) such that anc(*c*) is in the connected component and *c* outside, so the maximum of *t*(*c*) − *t*(anc(*c*)) over such *c* is the biggest negative value by which this connected component can move. Likewise *t*(anc(*r*(*s*_*i*_))) − *t*(*r*(*s*_*i*_)) is the maximum positive value by which this connected component can move. Thus any new times for the nodes *s*_*j*_ ∈ *C*(*s*_*i*_) that are in [*d*_*j*_, *u*_*j*_] for all *j* such that *s*_*j*_ ∈ *C*(*s*_*i*_) will preserve this connected component.

Put *δ* = *t′*(*S*) ≤ *t*(*S*), the amount by which the node times are changed. Then from step 6,

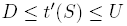

so

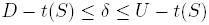

and for all *i*,

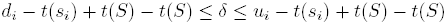

hence

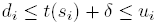

as required.

#### Proposition 2

The NodesNudge move is symmetric.

*Proof.* First note that the choice of *S* in step 1 has the same probability for the reverse move. Then, the key property of connected components described earlier together with Proposition 1, ensures that Δ(*S*) is unaffected by the move. It only remains to show that the interval [*D, U*] is unaffected by the move. Primes (*′*) are used to denote the various quantities after the move. From step 3, 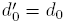 and 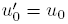, and for *i >* 0, from step 4 we have

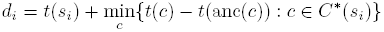

and since all the *t*(*c*) values are unaffected by the move, and all *t*(anc(*c*)) are changed by *δ*, as is *t*(*s*_*i*_), it follows that

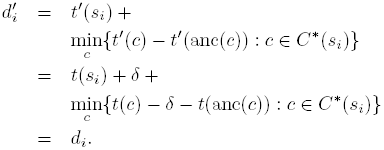

Similarly, 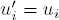 and it follows that [*D′, U′*] = [*D, U*]. Since the choice of *t′*(*S*) in [*D, U*] is uniform, the move is symmetric.

### 4.3 The definition of Δ(*S*)

In the NodesNudge move as currently implemented, Δ(*S*) is defined to be all the internal gene tree nodes *s* such that the pair (*S, s*) is hitched. For this definition of Δ(*S*), it is not possible for a node and its parent to both belong to Δ(*S*), so all the connected components *C*(*x*) in the algorithm consist of single nodes. This simplifies the implementation, and can be used to simplify the proofs of the two Propositions. However it is likely that future versions of STACEY will exploit the more general case.

## 5 The coordinated subtree and regraft move

### 5.1 The subtree prune and regraft move for one tree

First we describe the fixed height subtree prune and regraft algorithm [6] as it applies to a single tree. This is to make precise the algorithm used here, since there are variants of the main idea. Figure 3 illustrates the process. The algorithm prunes a subtree *S* and regrafts it into a branch *D* in both SMC-tree and gene trees. The requirements are that neither *S* nor anc(*S*) is the root of the tree, none of *S*, anc(*S*), or the sibling of *S* can be *D*, and *t*(*D*) ≤ *t*(anc(*S*)) ≤ *t*(anc(*D*)). It is possible for the *D* and anc(*S*) to be siblings (so in the figure, *Y* can be equal to *Z*). Note that there is no change in the set of node heights; the existing heights are re-used. The subtree prune and regraft algorithm for one tree follows.

**Figure 3:**
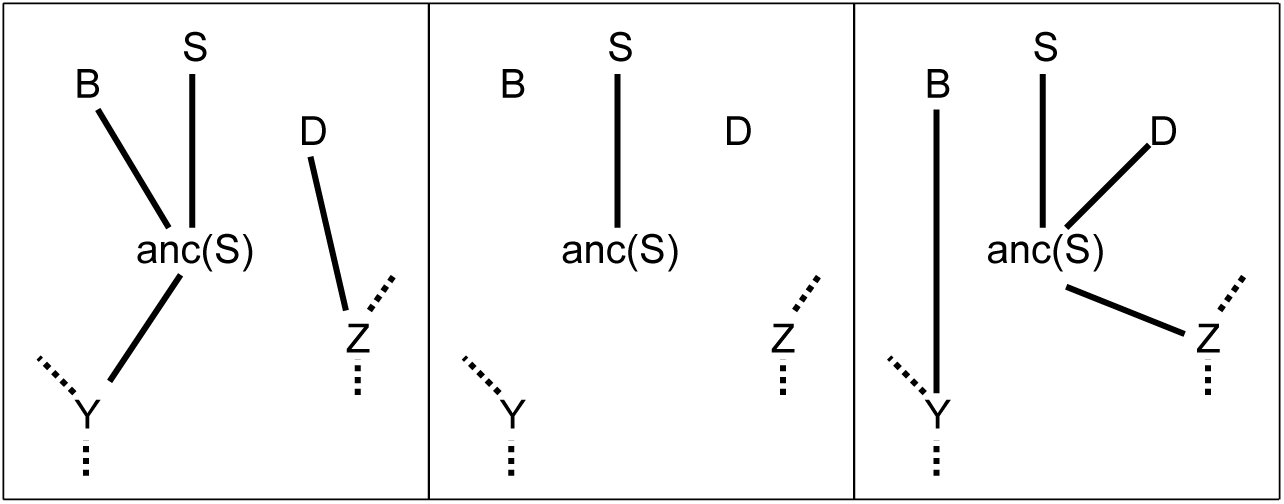
Fixed height subtree prune and regraft subroutine for one tree, shown in 3 stages from left to right.

1. Let *B* be the sibling of *S*, *Y* the parent of anc(*S*), and *Z* the parent of *D*.
2. Remove child *D* from *Z*. Remove child *B* from anc(*S*). Remove child anc(*S*) from *Y*.
3. Add anc(*S*) as child of *Z*. Add *D* as child of anc(*S*). Add *B* as child of *Y*.

### 5.2 The algorithm

The idea is to make a subtree prune and regraft move on the SMC-tree and a co-ordinated set of subtree prune and regraft moves on each of the gene trees in order to make them compatible with the new SMC-tree. Figure 4 shows an example. The node *S* is the subtree to be pruned and the branch *D* is the destination branch into which *S* is regrafted. Here is the algorithm.

**Figure 4:**
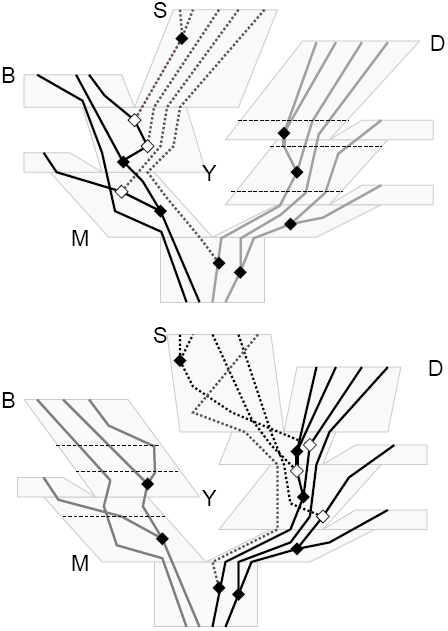
Example of the CoordinatedPruneRegraft move. The state before the move is at the top, and after the move below. The SMC-tree is pale gray. Gene tree branches whose sequences all belong to *I*(*S*) are dotted. Before the move, gene tree branches whose sequences are descendants of the same (left) child of *M* as *S* but do not belong entirely to *I*(*S*) are black. The white nodes are the origins (anc(s) in the text) of subtrees that need to be pruned and regrafted, and the thin horizontal dotted lines cut across the available destination branches. After the move, the colors and styles are reversed to illustrate the reverse move.

1. Choose at random any node *S* such that neither *S* nor anc(*S*) is the root. Let *B* be the sibling of *S*.
2. Choose at random any node *D* which is none of *S*, anc(*S*) or *B*, and such that *t*(*D*) *≤ t*(anc(*S*)) *≤ t*(anc(*D*)).
3. Find the most recent common ancestor node *M* of *S* and *D*.
4. For each gene tree *G*, find all the nodes *s* of *G* such that anc(*s*) is inside one the SMC-tree branches between the node anc(*S*) and the node *M* such that *I*(*s*) ⊂ *I*(*S*) and the sibling node *x* of *s* satisfies *I*(*x*) ⊄ *I*(*S*). Denote by Src(*G, S*) the set of such nodes *s*.
5. For each gene tree *G*, for each *s* ∈ Src(*G, S*), calculate the set of branches Dest(*G, s*) using the subroutine below.
6. Prune subtree *S* and regraft into branch *D*.
7. For each gene tree *G*, for each *s* ∈ Src(*G, S*), choose a member *d* of Dest(*G, s*) at random then carry out the prune and regraft operations for each pair.
8. For each transformed gene tree *G′*, let Src(*G′, S*) be defined as in step 4. For each *s′* ∈ Src(*G′, S*), calculate the set of branches Dest(*G′, s′*) using the subroutine below. Return

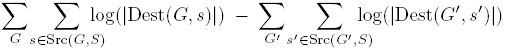

as the logarithm of the Hastings ratio.

In step 4, all the gene tree nodes which need to be pruned and regrafted are identified. These are nodes *s* all of whose sequences belong to *S*, and whose parent nodes anc(*s*) are the first nodes going back in time at which sequences from *S* are joined to those not belonging to *S*. In step 5, the set of possible destination branches is identified for each *s*. In steps 6 and 7, the prune and regraft operations are carried out, first for the SMC-tree, then all gene trees. In step 8, the number of choices for destination branches is found for the reverse move. The subroutine for the calculation of Dest(*G, s*) in steps 5 and 8 follows.

1. Set Dest(*G, s*) = *ø*.
2. Suppose the chain of nodes leading from *D* back to *M* is *D*_0_ = *D, D*_1_*, …, D_m_* = *M*. Let *x* be a node in *G*. Find *d* such that *t*(*D*_*d*_) *≤ t*(anc(*S*)) *≤ t*(*D*_*d*+1_)
3. For all nodes *x* in *G*, add *x* to Dest(*G, s*) if and only if *t*(*x*) *≤ t*(anc(*s*)) *≤ t*(anc(*x*)) and *I*(*x*) ⊂ *I*(*D*_*d*_).

Note that the number of choices in step 7 can differ for the reverse move. In the example of Figure 4, the numbers are 3,4,4 for the forward move, and 3,3,3 for the reverse move.

### 5.3 Properties

#### Proposition 3

The coordinated subtree and regraft move preserves compatibility

*Proof.* Pruning a subtree cannot produce an incompatibility, so we just need to show that the new nodes created by regrafting do not result in incompatibilities. The proof for each gene tree is identical, so we just consider one gene tree *G*. Suppose that the subtree *S* has been pruned and regrafted, and that all the nodes in Src(*G, S*) have been pruned but not yet regrafted. (The algorithm is not carried out in this order, but it is convenient for the proof.) The definition of Src(*G, S*) ensures that no incompatibility exists in this state. This is because the remaining nodes *x* in *G* either satisfy *I*(*x*) ∩ *I*(*S*) = *ø* or *t*(*x*) ≥ *t*(*M*).

It remains to consider the new nodes which are created when the gene subtrees are regrafted. Suppose node *x* is regrafted into branch *y*. The definition of destination branches Dest(*G, s*) ensures that *x* is created inside the branches between anc(*S*) and *M* and that *I*(*x*) only contains *I*(*y*) plus members of *I*(*S*). This shows that the new gene nodes are compatible, and completes the proof.

#### Proposition 4

The coordinated subtree and regraft move is reversible with Hastings ratio as given in step 8.

*Proof.* Given the source subtree and destination branch, each individual FNPR move is symmetric, so the only asymmetry arises in the choice of the set of moves. Since the move does not change heights, the number of branches whose duration includes a particular height *t* is unaffected by the move. It follows that the number of choices for *S* and *D* (steps 1 and 2) are the same for the reverse move: that is, the probability of choosing *D* for the forward move is the same as the probability of choosing *B* for the reverse move. Furthermore, it follows from the definition of Src(*G, s*) in step 4 that *|*Src(*G, S*)*|* = *|*Src(*G′, S*)*|*. Finally, the sizes of *|*Dest(*G, s*)*|* are accounted for in step 8.

## 6 The focused scaler move

### 6.1 The algorithm

We assume that the SMC-tree has at least 4 tips, so that the first step below is possible.

1. Choose at random any node *S* in the SMC-tree which is not the root or a tip, and has at least one child which is not a tip.
2. For any node *X* in the SMC-tree, let dist(*X*) be the number of branches from *S* to *X*.
3. For each gene tree *G*, find all the nodes *s* of *G* such that (*S, s*) are hitched.
4. For each node *s* hitched to *S*, define dist(*s*) to be 1 if *I*(*s*) ⊂ *I*(*S*) and 2 otherwise. For other nodes *x* in gene trees, dist(*x*) = ∞ initially. Then dist(*x*) is defined recursively using the following rule. If *y* is adjacent to *x* (that is, if *y* = anc(*x*) or *y* is a child of *x*), then dist(*x*) = min(dist(*x*), 1 + dist(*y*)).
5. For each tree (SMC-tree or gene tree) *T* let *f*_*T*_ : ℕ *→* [0, 1] be a function from the nonnegative integers such that *f*_*T*_ (0) = 1 and *f*_*T*_ (*d*) *<* 1 for *d >* 0. Let *w*(*X*) = *f*_*T*_ (dist(*X*)) for all nodes *X* of *T*.
6. Let Λ be the set of pairs of nodes defined as follows. Firstly, Λ contains all hitched pairs of nodes (*Y, y*) for any SMC-tree node *Y* and any node *y* in any gene tree. Secondly Λ contains all pairs (*X,* anc(*X*)) where *X* is in any tree. Thus Λ contains all hitched pairs and all branches.
7. Let Λ^+^ = {(*A, B*) ∈ Λ : *w*(A) > *w*(*B*)} and Λ^−^ = {(*A;B*) ∈ Λ : *w*(*A*) < *w*(*B*)}. Set

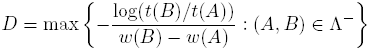

and

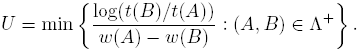
8. Choose *η* uniformly from the interval [*D, U*] and scale the height of every internal node *X* which has a nonzero weight by exp(*w*(*X*)*η*). Return the sum of all the *w*(*X*)*η* values as the logarithm of the Hastings ratio.

The conditions on *S* in step 1 ensure that there are two nodes adjacent to *S*, one with a bigger height (its parent) and one with a smaller but nonzero height (one of its children). Both these have distance 1 from *S* (step 2) so get a smaller weight than *S* due to the conditions *f*_*T*_ (0) = 1, *f* (1) *<* 1 in step 5. Thus the maximum and minimum in step 7 are taken over a non-empty sets, so *D* and *U* and hence *η* are all finite.

The nodes given a distance of 1 in step 4 are ‘topologically closer’ to *S* than those given distance 2. Usually, they are closer in height as well. There is freedom to choose a wide variety of functions for *f*_*T*_ in step 5. In the current implementation, a decreasing function is used, which becomes zero at the root of each tree. The weights *w*(*X*) are thus zero whenever dist(*X*) ≥ dist(*R*) where *R* is the root of the tree *T* containing *X*.

### 6.2 Properties

#### Proposition 5

The focused scaler move keeps all branch lengths nonnegative and all gene trees compatible with the SMC-tree.

*Proof.* Consider two nodes *X* and *Y* with (*X, Y*) ∈ Λ. Note that *t*(*X*) ≤ *t*(*Y*). Let *g*(*X*) = log(*t*(*X*)) and *g*(*Y*) = log(*t*(*Y*)). After the move the heights are *t*(*X*) exp(*w*(*X*)*η*) and *t*(*Y*) exp(*w*(*Y*)*η*) (step 8) so the logarithms of the heights after the move are *g′*(*X*) = *g*(*X*) + *w*(*X*)*η* and *g′*(*Y*) = *g*(*Y*) + *w*(*Y*)*η*. If *w*(*X*) = *w*(*Y*), we obviously have *g′*(*Y*) *≥ g′*(*X*). Suppose that *w*(*X*) > *w*(*Y*) so that (*X, Y*) ∈ Λ+. We have from step 7 that

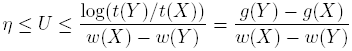

so

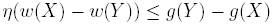

so the difference between the logarithms of heights after the move is

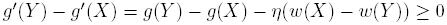

so it remains nonnegative. The case *w*(*X*) *< w*(*Y*) is similar.

Since Λ includes all branches in all trees (step 6), the move keeps all branch lengths nonnegative. It remains to show that compatibility is preserved. For pairs (*X, x*) ∈ Λ where *X* is a SMC-tree node and *x* a gene tree node, compatibility follows in the same way as for branch lengths. Suppose (*X, x*) is not in Λ. Then either *I*(*x*) does not straddle *X*, or at least one of *R*(*x*) or *L*(*x*) does straddle *X*. In the first case, *X* and *x* are compatible since there is no conflict between the minimal clusters belonging to them. In the second case, some descendant *y* of *x* must be in Λ, and the compatibility of (*X, y*) implies that of (*X, x*).

#### Proposition 6

The focused scaler move is reversible with Hastings ratio as stated in step 8.

*Proof.* The choice of *S* in step 1 is based on topological criteria only, and the move does not change the topology so the probabilities of choosing *S* for the move and reverse move are identical. The definition of dist() only depends on topologies and assignments at tips, so given the choice of *S*, the values dist(*X*) and hence *w*(*X*) for all nodes *X* are the same for the reverse move. Denoting quantities for the reverse move with primes (*′*) we have

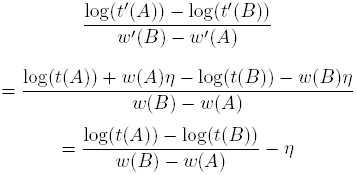

whenever *w*(*B*) ≠ *w*(*A*), from which it follows that

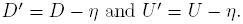

Now *D* ≤ 0 and *U ≥* 0, so 0 ∈ [*D, U*] and so *−η* ∈ [*D′, U′*], so a choice of *−η* is available for the reverse move, which will restore the original state. The two intervals [*D, U*] and [*D′, U′*] have the same size, and the choice of *η* is made uniformly, so there is no contribution to Hastings ratio here. This leaves only the scaling of the node heights for which step 8 provides the Hastings ratio.

## 7 Tests of correctness

In order to check the theory and the implementation of these moves, some tests were carried out by sampling from prior distributions. Full details of these tests and the results are in the supplementary information. The following is a brief summary. There were two sets of tests. One has an unknown number of species (between 1 and 8). The other uses a fixed number (8) of species and samples from the prior on the species tree. Although there is no sequence data, the assumptions about the number of species constitute some ‘meta-data’. In both sets of tests, there was one gene tree with no data, that is with a sequence “?” at each tip. Since the operators change the gene trees simultaneously with the SMC-tree, it is important to include at least one gene tree.

BEAST2 XML files were generated for the two sets of tests, with various combinations of operators. These were then run in BEAST2. The sampled SMC-trees were examined for agreement with theoretical distributions for the node heights; the species tree topology in the case of fixed species assignments; and for clusterings in the case of delimitation.

Some results in which the CoordinatedPruneRegraft move is used together with the usual BEAST operators are shown. These are included to illustrate the type of tests used, but for full details see the supplementary information.

Figure 5a is for the estimated delimitation case. It shows estimated and theoretical values for the 22 partitions of the number 8. Each partition of 8 represents one or more clusterings of 8 objects. There are a total of 4140 clusterings of 8 objects, and these can be grouped into 22 sets corresponding to the partitions of 8. For example suppose the 8 objects are *a, b, c, d, e, f, g, h*. One clustering is {{*a*; *b*; *c*}; {*d*}; {*e*; *f*; *g*; *h*}}, which corresponds to the partition 4+3+1 of 8. There are 280 clusterings with the shape 4+3+1, and whenever one of these are visited during the MCMC, it counts towards the posterior probability of this partition of 8.

**Figure 5:**
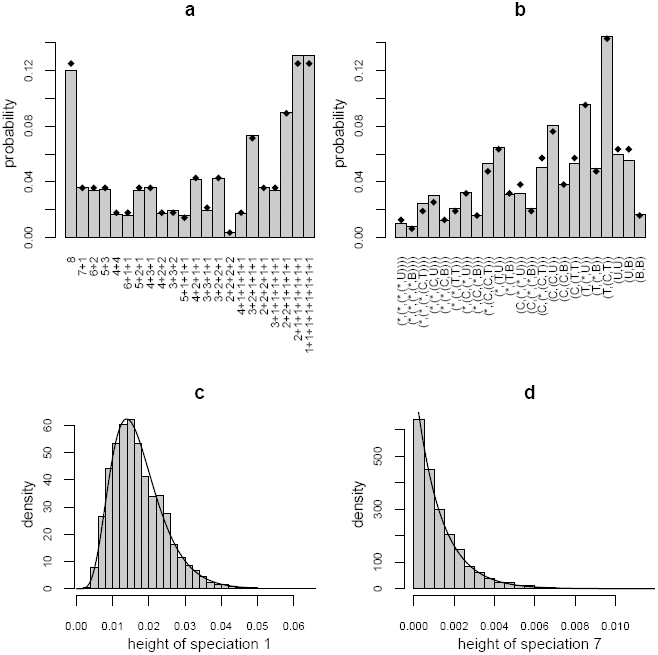
Plot (a) shows some results for the case of sampling from the prior distribution with an unknown number of species between 1 and 8. Estimated posterior probabilities are shown in the gray bars for the 22 partitions of the integer 8 (see text for further explanation). The theoretical values are shown as diamonds. Plots (b),(c), and (d) are some results for the case of sampling from the prior distribution with the number of species fixed at 8. In (b), the gray bars show estimated posterior probabilities for each of the 23 unlabeled rooted topologies. The theoretical values are shown as diamonds. The x-axis annotations enumerate the topologies (see text for details). In plots (c) and (d), the gray bars show a histogram of samples from the posterior for the first and 7th speciation height, that is, the root height and most recent speciation. The curves show the theoretical densities for these speciation heights.

The other three plots in Figure 5 are for the fixed species delimitation case. Figure 5b shows how this combination of operators sample from the 23 possible topologies with 8 tips. The x-axis annotations show the topologies in the following format. A tip is shown by **\***. A cherry **(*,*)** is denoted as **C**, a 3-tip tree **(*,(*,*))** as **T**, an unbalanced 4-tip tree as **U** and a balanced 4-tip tree as **B**. Otherwise, the Newick format is used. For example **(T,(*,U))** is short for **((*,(*,*)),(*,(*,(*,(*,*)))))**. Finally the sampled node heights are compared to theoretical densities in (5c,d).

The scenarios are sufficiently simple that various marginal aspects of the prior distribution can be calculated analytically, but sufficiently complicated to provide a meaningful test of the operators. No problems were found (after two bugs were found and fixed during preliminary testing). However, it is not possible to give a 100% guarantee of correctness. There may be problems which are not revealed by the marginal aspects of the distribution that were analyzed here. There may be problems which only produce an undetectable bias in these tests but which become more serious in other scenarios.

## 8 Results on the simulated data of Olave et al.

As a proof-of-concept demonstration, I re-analyzed the simulated data provided in the supplementary material of [10]. This data set contains 50 replicates for each of twelve configurations. In all cases there are 40 individuals, 2 sequences of length 1000bpp per individual, and 8 true species each consisting of 5 individuals. There are two tree shapes, symmetric and asymmetric, two amounts of incomplete lineage sorting, and three values 4, 8, or 14 for the number of loci.

The data was incorporated into XML files for BEAST2. Version 2.2.0 of BEAST2 and 1.0.1 of STACEY were used. For each replicate, the program was run for 5, 7, or 10 million generations for the 4, 8, and 14 loci cases respectively. The first 1 million discarded as burnin. Samples were taken every 1000 generations, so there were 4000, 6000, or 9000 SMC-trees on which to base the species delimitations using SpeciesDelimitationAnalyser [9].

### 8.1 Priors and other settings

For the population variability among branches, a single inverse gamma component with mean and standard deviation 1 was used. (In equation (3), *C* = 1*, α*_1_ = 3.0*, β*_1_ = 2.0.) A lognormal(-7.0,2.0) was used for the hyperprior *π*_*σ*_ for the overall population scaling factor. (Parameters to the lognormal are given in log space.) The value of *p*_*j*_ was set to 2 for all genes. The HKY model was assumed for the substitution model. It was assumed that there was no site rate heterogeneity (although the data set does contain such heterogeneity). The relative clock rates of the genes other than the first were estimated; a lognormal(0.0,1.0) prior was assumed for these. A birth-death model was assumed for the species tree, with a lognormal(4.6,2) hyperprior for the growth rate, and a Beta(3,1) hyperprior for the relative death rate. The prior on the collapse weight was uniform on [0, 1] so that there was a flat prior on the number of species, and the collapse height *∈* was set to 0.0001. The 40 individuals were used as minimal clusters (containing two sequences each) in STACEY. (See [9] for definitions of ‘minimal cluster’, ‘collapse weight’ and ‘collapse height’.)

### 8.2 Results

The results are shown in Figure 6. The clustering with the largest posterior probability (that is, a MAP estimator) was used to estimate the species delimitation. All errors in this estimate were false splits. Usually just one of the true species was split; in five replicates, two true species were split; and in replicate 47 from YH4 and replicate 34 from ZH4, three true species were split. In all 600 replicates, the true clustering was in the 0.95 credible set. The highest posterior probability assigned to a erroneous clustering was 0.83 (replicate 16 from YE4).

The estimated sample sizes (ESSs) for the posterior, as reported by Coda [11], had means of 250, 215, and 215 for the the 4, 8, and 14 loci cases. Some individual replicates had ESSs below 100, with a minimum of 72 over all 600 replicates.

## 9 Discussion

Based on tests so far, including some results not reported here, STACEY converges much faster than DISSECT. The difference was particularly apparent in the time to ‘burn-in’ in cases where there was little ILS and many loci. Although these are the cases where signal is strongest, DISSECT can take a very long time to converge, as reported in [9] for the case of 27 loci. It is not yet clear how much of this improvement is due to the new model and how much to the new moves. It seems likely that the new moves will improve convergence in *BEAST in some cases at least, but this has not been tried.

When analyzing the data [10], the number of generations in the MCMC chains were chosen so that all 600 replicates could be run in a reasonable amount of time with limited computational resources (2 weeks on a desktop computer with 4 cores). This resulted in lower ESS values than desirable on some replicates. Given the main purpose of this analysis, this does not seem important: if anything longer runs would be expected to improve accuracy. When used ‘for real’, several longer runs are strongly recommended.

In the context of phylogeny estimation, the relative importance of the two kinds of noise, namely mutational variance and incomplete lineage sorting, was studied in [7]. In their scenarios, up to 75% of the errors in maximum likelihood estimates of species trees were attributable to mutational variance. It seems very likely that similar conclusions apply to Bayesian species delimitation. The simulated data sets of [10] have low mutational variance. The species tree branch lengths, measured in substitutions, range from 0.004 to 0.028 in the N=0.4 case and from 0.04 to 0.28 in the N=4 case. Since there are two sequences of length 1000bp per individual, the expected number of substitutions per individual per locus along a branch is always at least 0.004 * 2000 = 8. However, in many empirical data sets the difficulties due to incomplete lineage sorting will be compounded with large amounts of mutational variance. The simulations used in [9] were much harder in terms of the mutational variance: the sequences were 500bp, there was only one sequence per individual, and the shortest branch lengths were 0.001, so that the expected number of substitutions along the shortest branches is only 0.5 instead of 8. The results of that paper may be a better guide to to the accuracy of the approach on many empirical data sets.

**Figure 6:**
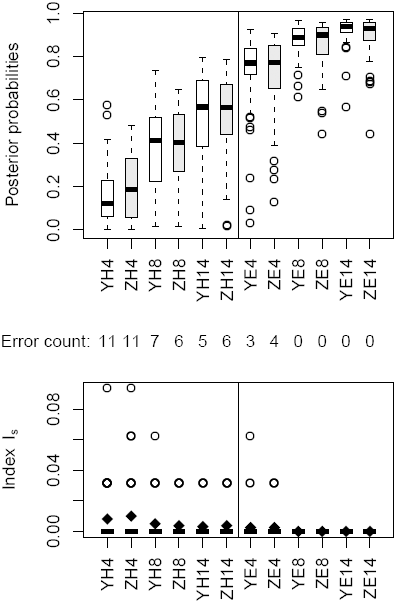
The upper boxplots show the posterior probabilities of the true clustering over 50 replicates for twelve configurations YH4,… ZE14. In the labels for the configurations, the first letter Y or Z denotes the tree shape, with Y for symmetric and Z for asymmetric; the second letter denotes the degree of incomplete lineage sorting, with H for N=0.4 (‘hard’) and E for N=4 (‘easy’); this is followed by the number of loci: 4, 8, or 14. The numbers between the boxplots are the number of times out of 50 that the clustering with the largest posterior probability was not the true clustering. The lower boxplots show the measure of over-splitting of true species lineages using the index *I*_*s*_ of Olave et al (2014). The black diamonds show the mean values. Note that the vertical scale is much smaller than that of Figure 3 in Olave et al.

The results here should dispel some of the pessimism expressed in [10] about DNA-based species delimitation. It is usually the case that geographical and morphological information is available as well [18], but it is rare that this provides certainty about the assignment of individuals to clusters or populations. I think that a more promising way ahead is to include the extra information in a Bayesian analysis. The location data and morphological characters could be included alongside the genetic data. Alternatively, taxonomists could formalize their knowledge in the form of a prior on the space of all possible clusterings. A program like STACEY can then explore the full space, taking into account the extra information. The space of all clusterings is huge, and it is not easy to construct sensible probability distributions for it which reflect expert knowledge about the organisms. Research is needed to find good ways of doing this.

## Acknowledgments

I thank the developers of BEAST for making this work feasible, and Remco Bouckaert in particular for helpful advice on writing the STACEY package. I thank the authors of [10] for making their simulated data readily available, and Melisa Olave for supplying extra details about the simulations.

